# Structural integrity following focused ultrasound thalamotomy and its correlation with tremor relief

**DOI:** 10.1101/620369

**Authors:** Gil Zur, Orit Lesman-Segev, Ilana Schlesinger, Dorith Goldsher, Alon Sinai, Menashe Zaaroor, Yaniv Assaf, Ayelet Eran, Itamar Kahn

**Author notes:** These authors contributed equally to this work (Gil Zur, Orit Lesman-Segev).

## Abstract

**Background:** Magnetic-resonance-guided focused ultrasound ablation of ventral intermediate nucleus of the thalamus is a new treatment for tremor disorders.

**Objectives:** We sought to evaluate the white matter integrity prior to and following focused ultrasound ablation and its correlation with clinical outcome.

**Methods:** 22 patients with essential tremor and 17 patients with Parkinson’s disease underwent tremor and quality-of-life assessments prior to and at one and six months following focused ultrasound ablation. All patients underwent MRI including T1, T2-FLAIR and diffusion weighted imaging before treatment and at one day, 7–10 days, 1–3 months, and 6 months or more following it. Diffusivity parameters were calculated and fiber tractography measures were extracted. Change in diffusivity parameters were assessed inside the ablated core, in the motor thalamus and throughout the dentato-rubro-thalamo-cortical tract.

**Results:** We found short-term changes in the motor thalamus and in the tract between the thalamus and regions within the dentato-rubro-thalamo-cortical tract. Long term damage was found in the ablated core and in the tract connecting the thalamus and red-nucleus. Inverse correlation was found between fractional anisotropy in the motor thalamus one day following ablation and tremor improvement in both patient groups, with lower values before treatment associated with better outcome (tremor relief) in essential tremor patients.

**Conclusions:** Short-and long-term changes in white matter integrity are present following focused ultrasound thalamotomy. Regions demonstrating long-term white matter changes may be responsible for the tremor relief seen in patients, implicating these regions in the disorder’s pathogenesis.

## Introduction

Essential tremor and Parkinson’s disease are the most prevalent tremor disorders.^1,2^ Essential tremor is characterized by upper limb intention or postural tremor, and is considered mostly a “pure” tremor disease compared to Parkinson’s disease, which is characterized by a variety of clinical manifestations, amongst them, rest tremor, which is seen in about 75% of the patients.^3^ Both diseases have a significant influence on quality of life,^4,5^ and are associated with a heavy economic burden to both patients and society.^6^

A number of hypotheses exist regarding the pathogenesis of these diseases. Neurodegeneration of the cerebellar dentate nucleus, abnormal electrical rhythmicity and reduced function of the inhibitory neurotransmitter GABA are possible mechanistic explanations for essential tremor.^7^ Tremor in Parkinson’s disease is related indirectly to dopamine deficiency and responds only partially to dopamine treatment. Therefore, the pathogenesis of tremor in Parkinson’s disease is suggested to differ in its underlying cause relative to other Parkinson’s disease symptoms, such as rigidity and bradykinesia.^8^ Alternatively, it has been proposed that disruption in basal ganglia dopaminergic neurons results in imbalance and activation of the tremor network.^7^ Consequently, it is possible that the dentato-rubro-thalamo-cortical (DRTC) pathway is a common component in tremor pathogenesis.^7,9^

The DRTC pathway consists of cerebellar efferent fibers of the contra-lateral thalamus and red nucleus, and projections from the thalamus to the motor cortex. One of the structures interacting with the DRTC pathway is the Guillain-Mollaret triangle, which includes connections between the inferior olive, dentate nucleus and red nucleus, and forms a feedback mechanism for the cerebellum.^10^ Although the mechanism of damage to the DRTC pathway is unclear, the potential involvement of white matter tracts in tremor pathogenesis highlights the importance of brain imaging methods such as diffusion tensor imaging (DTI) in assessing white matter integrity in tremor patients. Previous studies of DTI in essential tremor patients showed reduced fractional anisotropy (FA), which may reflect damage to white matter integrity in the superior and inferior cerebellar peduncle (ICP), dentate nucleus, cerebellar cortex, pons and other areas outside the DRTC tract.^11–13^ Additional studies reported increased mean diffusivity (MD) in ICP,^11^ and increased apparent diffusion coefficient (ADC) values in the red nucleus,^14^ both measures of white matter integrity. Reduced FA was reported in mild to moderate Parkinson’s disease patients in the substantia nigra, basal ganglia and several cerebral cortical areas.^15,16^ Finally, a recent study demonstrated reduced FA along the DRTC tract following MRI-guided focused ultrasound (MRgFUS) ablation of the thalamic ventral intermediate nucleus (VIM) in essential tremor patients.^17^

Thermal ablation of VIM using MRgFUS has emerged in recent years as a promising minimally invasive procedure with a high success rate for medication refractory tremor in essential tremor patients,^18–20^ and for medication refractory rest and action tremor in Parkinson’s disease patients,^21^ In this procedure, MRI-based anatomical landmarks are used to locate the thalamic VIM, followed by heating the nucleus using ultrasound energy (sonications). Temperature is increased gradually until there is a reduction in the tremor with no side effects, at which point temperature is increased to induce permanent ablation. As the VIM is arranged according to somatotopic representation, the procedure can target specific areas. Thus, the hand area is treated first since it is most disabling, followed by other areas (head, chin, leg, etc.) as needed. The most common reported adverse events are paresthesia and gait disturbance.^18–20^

By using longitudinal imaging of white matter integrity with diffusion-weighted imaging pre-and post-ablation, we aimed at characterizing short-and long-term white matter changes, and at using these changes to better understand tremor pathogenesis and identify factors contributing to treatment success. We hypothesized that tremor improvement is correlated with white matter changes in the DRTC tract and that characterization of white matter properties would aid in patient selection for thalamotomy.

## Materials and Methods

### Study Participants

The study included 39 participants (22 essential tremor, 17 Parkinson’s disease). Summary group statistics are reported in **Table 1**. Patients with essential tremor or Parkinson’s disease diagnosed by a neurologist specializing in movement disorders were enrolled on the basis of eligibility criteria as previously described.^18,19^ Average age (±SD) in the essential tremor group was 71.9±6 years, and 65.2±7.9 years in the Parkinson’s disease group. There was a male preponderance in both groups (64% of the essential tremor patients and 76% of the Parkinson’s disease patients were males). Most patients were right-handed (essential tremor: 73%; Parkinson’s disease: 88%). Average disease duration was 13.1±8.3 years in essential tremor patients and 6.6±3.4 in Parkinson’s disease patients. Written informed consent was obtained from all the patients included in the study.

**Table 1.** Baseline demographics and clinical characteristics of the study participants

|  |  |  |  |
| --- | --- | --- | --- |
|  | **ET (N = 22)**<br>**Discovery** | **ET (N = 17)**<br>**Replication** | **PD (N = 17)** |
| Age — year | 71.91 ± 6.01* | 73.47 ± 6.32 | 65.24 ± 7.86 |
| Male sex — no. (%) | 14 (64) | 10 (59) | 13 (76) |
| Duration of tremor — year | 13.12 ± 8.28 | 12.29 ± 9.35 | 6.62 ± 3.43 |
| Right handed — no. (%) | 16 (73) | 15 (88) | 15 (88) |
| Right hand treatment — no. (%) | 16 (73) | 15 (88) | 14 (82) |
| Tremor ‡ |  |  |  |
| pre ablation | 19.24 ± 6.85 | 18.24 ± 4.25 | 5.35 ± 1.58 |
| 1 month post ablation | 3.14 ± 3.66 | 2.94 ± 3.54 | 1.5 ± 2.13 |
| 6 months post ablation | 4 ± 4.04 |  | 0.8 ± 2.20 |
| Quality of life § |  |  |  |
| pre ablation | 47.15 ± 18.39 | 40.59 ± 20.99 | 43.82 ± 20.94 |
| 1 month post ablation | 14.05 ± 19.87 | 17.29 ± 16.87 | 27.07 ± 20.37 |
| 6 months post ablation | 13.38 ± 6.88 |  | 25 ± 24.17 |

\* Mean ± SD
‡ For essential tremor, the tremor score was derived from Clinical Rating Scale, which ranges from 0 to 32 for tremor in the hand. For Parkinson's disease, the tremor score was derived from Unified Parkinson's Disease Rating Scale which ranges from 0 to 8 for action and rest tremor in the hand
§ For essential tremor, quality of life was measured by the Quality of Life in Essential Tremor (QUEST) self-reported questionnaire which ranges from 0 to 120. For Parkinson's disease, quality of life was measured by the Parkinson's Disease Questionnaire (PDQ-39) which ranges from 0 to 156

### Clinical Outcome Assessment

Tremor scores were assessed before treatment and at follow-up visits (one and six months after treatment). For essential tremor, tremor assessment of the treated hand was based on the tremor part of the Clinical Rating Scale for Tremor (CRST).^22^ In this scale, the score for each item is 0 (no tremor) to 4 (severe tremor), with a maximal score of 32. For Parkinson’s disease, tremor was measured at rest and in action by a sum of questions 20 and 21, respectively, in the motor part (part III) of the Unified Parkinson’s Disease Rating Scale (UPDRS).^23^ In this scale, the score for each item is 0 (normal) to 4 (severe abnormal). Change in tremor score was defined as the relative reduction in tremor score: 100 × [(tremor score at baseline) – (tremor score at 1 month)] ÷ (maximal tremor scale score). Quality of life in patients with essential tremor was measured by the Quality of Life in Essential Tremor (QUEST) self-reported questionnaire.^24^ Quality of life in patients with Parkinson’s disease was assessed by the Parkinson’s Disease Questionnaire (PDQ-39).^25^

### Data Acquisition

All MR imaging studies were performed using the same 3T system (Discovery™ MR750, GE Healthcare, Chicago, IL, USA). Patients were recruited between 2016– 2018 and scanned prior to the procedure and in four follow-up scans. The first post-treatment scanning timing was identical across participants, but the next time points varied: 8.5±1.67 (6–14), 55.4±18.91 (27–101), and 344.7±146.44 (136–81) days (average±std [minimum–maximum]). We termed the five scanning time points: Baseline, 1 Day, 7–10 Days, 1–3 Months, and 6+ Months. Three-dimensional spoiled gradient-recalled echo (SPGR), axial T1-weighted images, diffusion tensor imaging (DTI) and T2-weighted fast fluid attenuated image recovery (FLAIR) were acquired in this order within the scanning session at each time point. T1-weighted 3D-SPGR was acquired with the following parameters: whole-brain coverage, TR/TE = 8.3/3.23 ms, FA = 12°, resolution 0.86×0.86×0.6 mm^3^. Eight minutes of DTI was acquired with the following parameters: whole-brain resolution 2×2×2 mm^3^, TR/TE = 9538/82 ms, FA = 90°, Δ/δ = 33/26, b = 1000 s/mm^2^, with 25 gradient directions and an additional five images with no diffusion weighting (B0 image). T2-weighted FLAIR was acquired with the following parameters: TR/TE = 8779/127 ms, FA = 111°, resolution 0.43×0.43×5 mm^3^. Participant withdrawal limited imaging data contribution to the different time points (1 Day: n = 39/39/38 [DTI/T1/T2-FLAIR]; 7– 10 Days: 35/35/34; 1–3 Months: 27/27/26; 6+ Months: 25/25/25).

### Image Analysis

ANTS (Advanced Normalization Tools) were used to non-linear register the T1-and T2-weighted scans to the Montreal Neurological Institute (MNI) atlas. Affine registration was used to register each diffusion-based scan to its corresponding T1-weighted acquisition. These transformations were concatenated to produce one transformation from the diffusion-based scans to the MNI atlas. The ablated lesions were identified one day post-ablation by the signal intensity threshold below 2000 for T1-weighted imaging. In order to capture both cytotoxic and vasogenic edema one day post-ablation, we used signal intensity threshold above 900 for T2-FLAIR imaging. These thresholds were estimated by subtracting for each participant the histogram of signal distribution of scans post-procedure from the pre-procedure scans. An expert radiologist evaluated each set of brain scans for accuracy of lesion and edema volume segmentation. Center-of-gravity was calculated for each lesion to find the maximal change in FA. The edema has irregular shape and therefore center-of-gravity doesn’t represent the true center. A probabilistic atlas,^26^ was used to parcellate the thalamus into six different segments and define seven ROIs along the tremor tract: dentate nucleus, superior cerebellar peduncle, cerebral peduncle, red nucleus, motor thalamus, posterior limb of internal capsule, superior corona radiata and precentral gyrus.

### Diffusion-weighted Imaging Analysis and Tractography

Diffusion-weighted images were analyzed using ExploreDTI.^27^ Images were first regularized and resampled to a resolution of 2×2×2 mm^3^ using a B-cubic spline fitting algorithm. Next, tensors were calculated using a robust estimation algorithm,^20^ and images were corrected for motion distortions and eddy currents. Corrections for field inhomogeneity-induced distortions were executed based on T1 images. No significant differences in rotation, translation and scaling were found along the different time points (**Supplementary Fig. 1**). Maps of mean diffusivity (MD), fractional anisotropy (FA), radial diffusivity (RD), axial diffusivity (AD) and B0 images were extracted.

**Figure 1:**
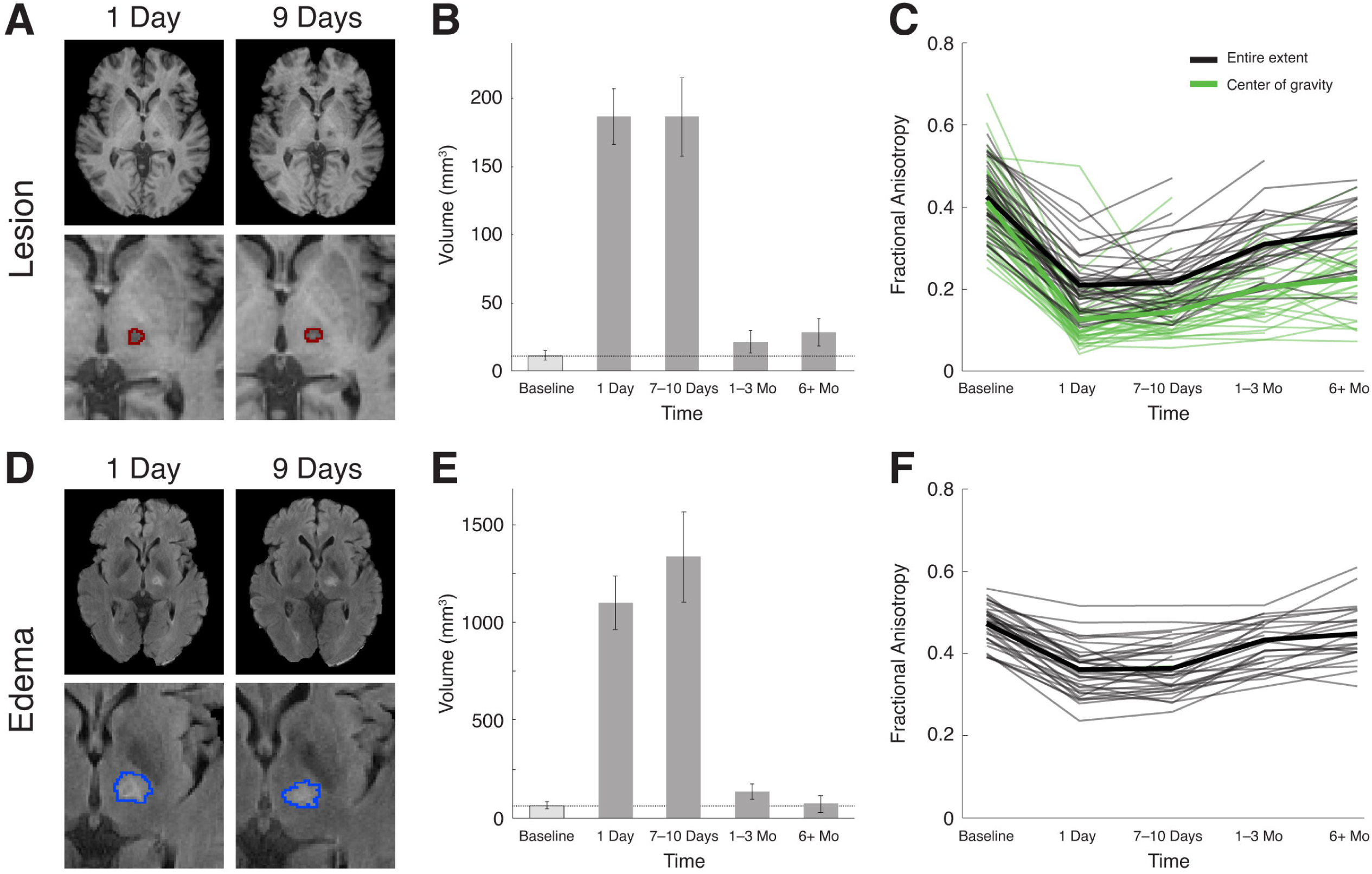
Lesion (A-C) and surrounding edema (D-F) following focused ultrasound ablation (FUS) of the thalamic ventral intermediate nucleus (VIM). The lesion and surrounding edema demonstrate maximal volume during the first 10 days following ablation, with a small remnant lesion though full recovery of the edema after 6 or more months. **(A)** Representative images at the level of thalamus, demonstrating the lesion extent (MRI T1) at one and nine days following ablation. **(B)** Mean volume (±SEM) of the lesion at one day, 7–10 Days, 1–3 Months and 6+ Months following ablation compared to the baseline value that represents the measurement error. **(C)** Fractional anisotropy (FA) in the area of the lesion at all follow-up time points. **(D)** Edema extent (T2 FLAIR), **(E)** mean volume following ablation and **(F)** FA are depicted. In **(C)** and **(F)**, thin lines represent FA values of individual participants and thick lines represent group mean for the entire extent (*black*) or center of gravity (*green*) of the lesion or edema.

Fiber orientation distribution (FOD) was reconstructed using the constrained spherical deconvolution (CSD) method in ExploreDTI. Whole-brain FOD-based tractography was performed in native and corrected spaces for all participants. Spherical harmonics up to the sixth order were used in the estimation. An FOD threshold of 0.1, a maximum angle deviation of 30 degrees and a step size of 1 mm were used. The minimum length of the fiber was set to 50 mm and the maximum to 500 mm.^27^ For each tract, number of tracts (representing an estimation of the number of fibers in the entire tract) and tract volume (number of voxels in the tract) were calculated. To select the tracts of interest from the whole-brain tractography, regions of interest (ROIs) were defined based on *a priori* information of tract location. In particular, the “tremor tract” was calculated between the red nucleus and subcortical WM.^28^ We defined the tract based on this approach because using the dentate nucleus as a seed showed unreasonable tractography results (see Limitations section below). MD and FA were calculated for each of the 50 equal-length segments of the tract. Inter-regional connectivity was then examined by determining the number-of-tracts (NOT), tract-volume (TV) and FA between the motor thalamus and the different ROIs. For the DRTC tract, we calculated the tract density (i.e. number of tracts per each voxel), FA and MD as well.

### Statistical Analysis

Statistical analysis was mainly done to assess the difference between values in each follow-up scan compared to the pre-ablation value. Paired student *t*-tests were used to assess changes in lesion and edema volumes and FA given a normal distribution of the data. NOT and TV tractography measurements were evaluated using the non-parametric Wilcoxon signed-rank paired two-tailed test since these data exhibited non-normal distributions. Pearson correlation values were calculated between the FA values, pre-and one day post-ablation, to the clinical outcome.

## Results

### Impact of Ablation on the Thalamus

The evolution of the lesion and surrounding cytotoxic and vasogenic edema and its effect on the DRTC tract were evaluated using structural and water diffusivity measures. Lesion volume was estimated by segmenting T1-weighted MRI scans for each time point, and structural changes in the lesion itself were evaluated using FA (**Fig. 1A-C;** see **Supplementary Fig. 2** for an example of the temporal changes following ablation according to ADC, FA, T1 and T2 maps, with the calculated tract between the red nucleus and precentral gyrus). Maximal lesion volume was observed at One day and 7–10 Days following ablation, exhibiting a dramatic decrease in size within the first couple of months, with only a minimal lesion observed at later follow-up time points (**Fig. 1B** and **Supplementary Table 1)**. Two different estimates of FA were used to evaluate changes in structural integrity: the entire extent of the lesion, and the voxel at which the volume of interest is evenly dispersed, and all edges are in balance (center of gravity). FA values significantly decreased within the entire extent of the lesion and in the center of gravity (**Fig. 1C**), reaching a negative peak one day after ablation. Incomplete recovery in FA was seen 7–10 Days to 6+ Months following the procedure. Recovery was poorest in the center of the lesion. All participants showed a decrease in FA followed by partial or complete recovery. While most patients reached the lowest FA values one day after ablation, some reached their lowest values 7–10 Days after ablation. FA values were reliably smaller across both measures relative to the pre-ablation period. As expected, edema covered a much larger volume compared to the lesion (**Fig. 1D-F** and **Supplementary Table 1**), though it followed a similar spatiotemporal evolution with only minimal edema observed one month to three months following the procedure and none at 6+ Months both in terms of volume (**Fig. 1E**) and decreased FA values (**Fig. 1F**).

**Figure 2:**
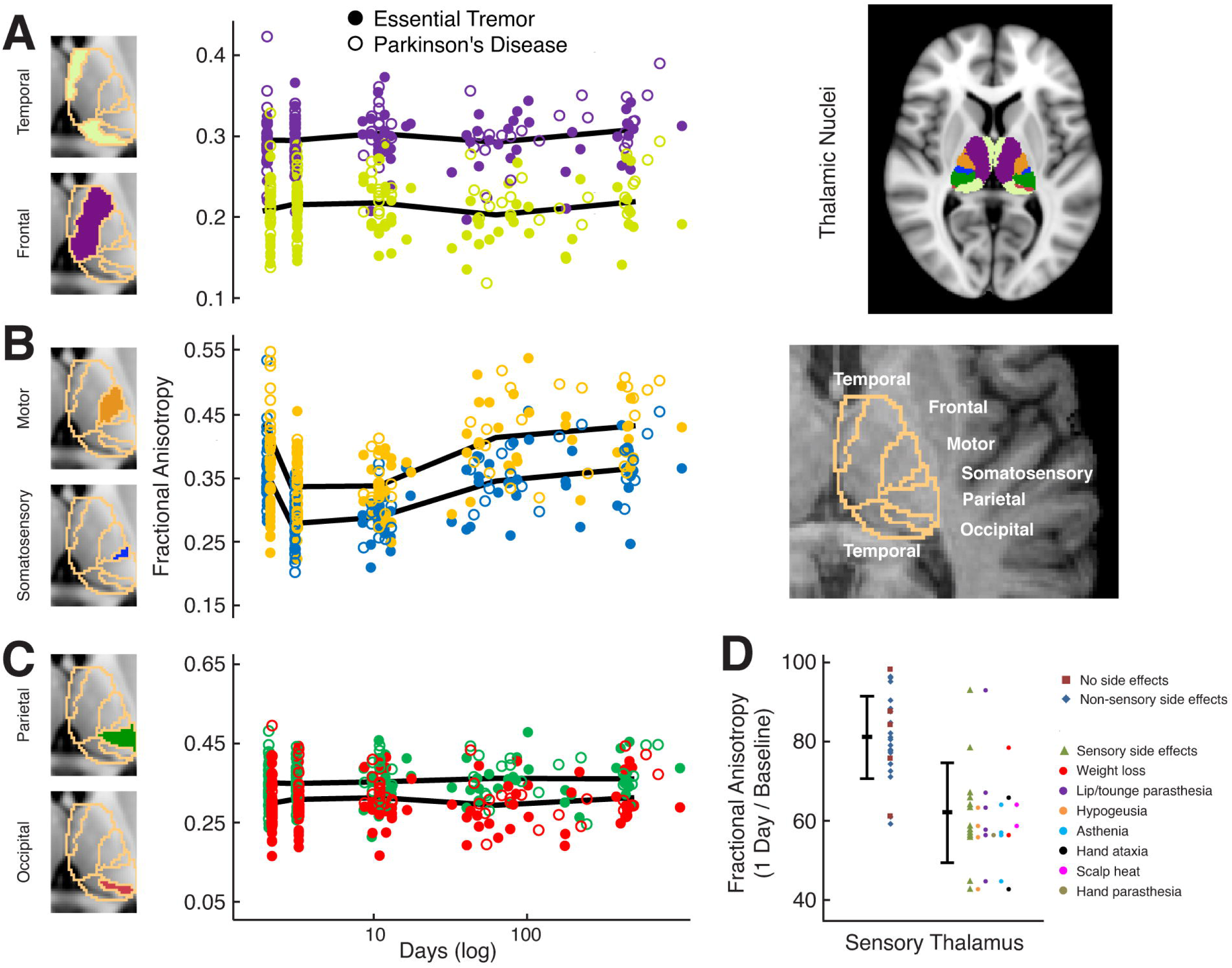
Ablation of the motor nucleus results in a localized and temporally restricted effect on the measures of white matter integrity. Parcellation of the thalamic nuclei derived from the Montreal Neurological Institute (MNI) atlas shows subdivisions into temporal, frontal, motor, somatosensory, parietal and occipital regions. **(A)** Fractional anisotropy (FA) within the temporal and frontal subdivisions demonstrates negligible impact of the ablation procedure. **(B)** Motor and somatosensory subdivisions demonstrate a decrease in FA values immediately following the ablation procedure and recovery after 6 or more months. **(C)** Parietal and occipital subdivisions show negligible impact of the ablation procedure. Study participants (*dot*) and mean FA (*line*) are depicted. **(D)** FA percent change in sensory thalamus one day post ablation relative to baseline. Participants with sensory side effects have a greater reduction in FA relative to participants with no side effects or side effects not categorized as sensory.

Next, we sought to evaluate the impact beyond the lesion and edema regions of the ablation procedure in the thalamus. We evaluated changes in FA across the temporal, frontal, motor, somatosensory, parietal and occipital subdivisions of the thalamus as defined by the probabilistic atlas (**Fig. 2**). A decrease in average FA values was noted only in the motor and sensory thalamic subdivisions, which were followed by near complete recovery in both areas, approaching pre-ablation values by the follow-up after 6 or more months (see **Supplementary Table 2**). Comparing the FA values after treatment to those before treatment revealed reliable differences only in the sensory and motor areas (**Fig. 2B**). All the other thalamic subdivisions (temporal, frontal, parietal and occipital) showed minor changes in FA values throughout the follow-up period relative to FA values before treatment (**Fig. 2A,C**). Finally, FA percent change in sensory thalamus one day post ablation relative to baseline were evaluated in respect to side effects (**Fig. 2D**). Participants with sensory side effects demonstrated a greater reduction in FA relative to participants with no side effects or side effects not categorized as sensory (*P*<0.001, t_37_=2.03).

### Impact of Ablation on the Dentato-rubro-thalamo-cortical Tract

The impact of ablation on the implicated fiber tract was evaluated next. The cerebral peduncle, precentral gyrus white matter, superior cerebellar peduncle and red nucleus, as well as the posterior limb of the internal capsule and superior corona radiata, which comprise the DRTC tract, were evaluated. FA values did not demonstrate a reliable change along any of the fiber bundles. In contrast, the number of tracts did demonstrate a reliable decrease in all selected ROIs along the DRTC tract one day after ablation (**Fig. 3A**). This decrease was followed by gradual and complete recovery in all ROIs, with the exception of the red nucleus, which showed only partial recovery in these parameters throughout the follow-up period. Similar values as well as patterns of decrease followed by complete recovery were seen in all ROIs in both essential tremor and Parkinson’s disease patients (representative ROIs are shown in **Fig. 3B**; see **Supplementary Table 3** for data and statistics in both groups and for both tract volume and number of fibers).

**Figure 3:**
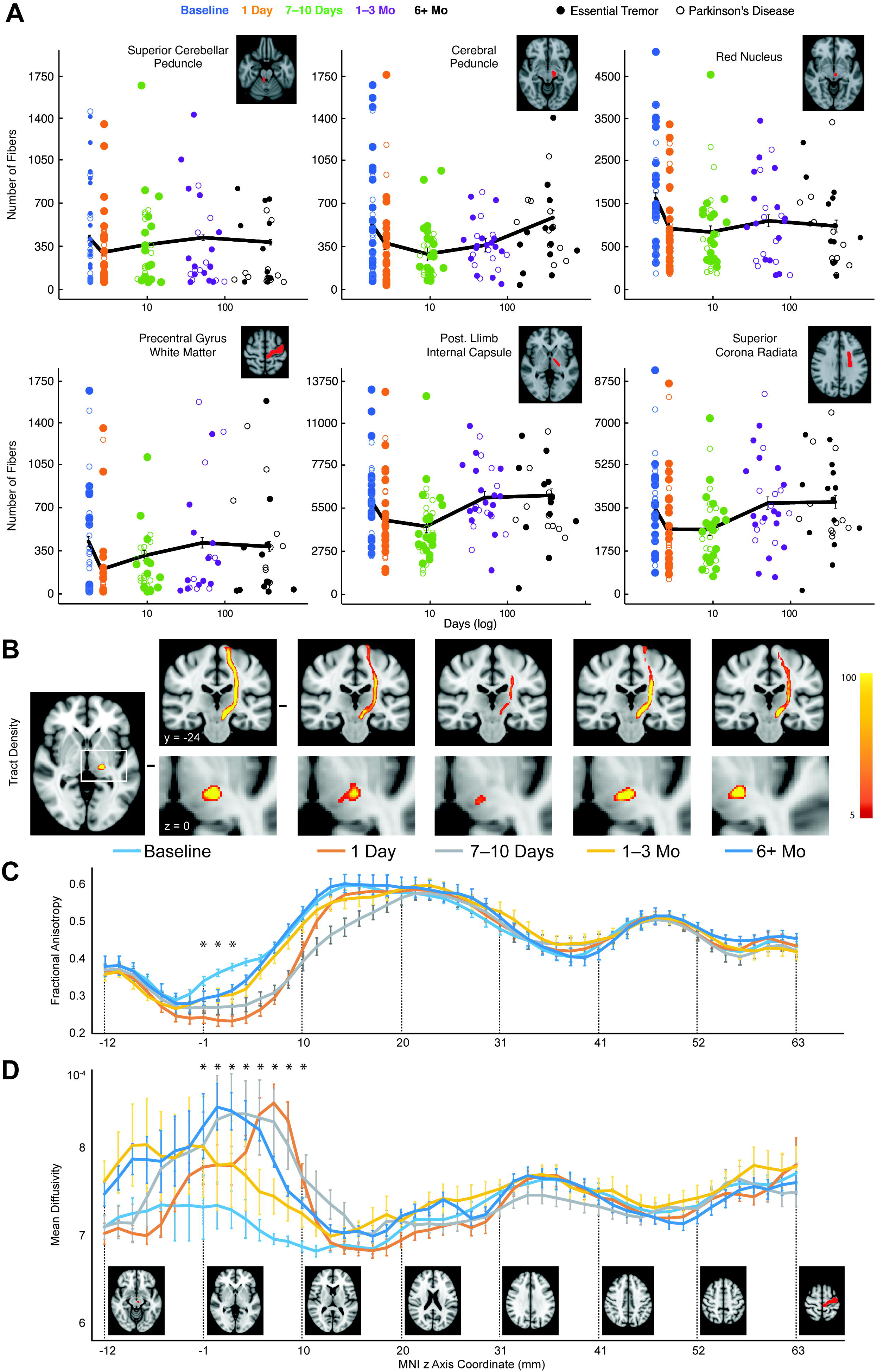
Measures of central white matter axon fiber bundles integrity demonstrate short-and long-lasting impact of ablation in both essential tremor (ET) and Parkinson’s disease (PD) study participants. **(A, B)** Six regions implicated in tremor were defined: Superior Cerebellar Peduncle (SCP, contralateral to the ablation site), Cerebral Peduncle (CP), Red Nucleus (RN), Posterior Limb of Internal Capsule (PLIC), Superior Corona Radiata (SCR) and Precentral Gyrus white matter (PCG). Region of interest extent (*red*) is depicted on a representative transverse T1 MRI. **(A)** Number of fibers are depicted for SCP, CP, RN and PCG, demonstrating a short-lasting decrease in the number of fibers for CP, PCG and SCP, and long-lasting impact on RN. **(B)** Number of fibers were calculated separately for ET and PD participants in PLIC and SCR (and in all the other regions; not shown), demonstrating short-lasting effect in both groups. **(C)** As can be appreciated qualitatively, measures of axon fiber integrity between the Red Nucleus and Precentral Gyrus white matter demonstrate a spatially restricted long-lasting impact of the ablation procedure and temporally restricted impact in regions around the ablation focus. Tract density is shown prior to and after the ablation procedure in the coronal at *y* = −24 (*top*) and axial *z* = 0 (*bottom*) slices depicting the thalamus. **(D, E)** Quantification of this effect is done by measures of fractional anisotropy (FA) and mean diffusivity (MD). Plotted from inferior to superior along the tract (z = 0 represents the ablation site), temporally long-lasting impact is revealed in both measures in *z* at around −1 to 6, and short-lasting temporal impact of ablation in FA (but not MD) in z at around −3 to 19.

Lower tract density was seen in the entire tract as of the first day following ablation, reaching a negative peak one week following ablation, with recovery later in the follow-up period (**Fig. 3C**). Changes in FA and MD throughout the tract were more extensive locally (closest to the ablation site) than far away from the ablation site (**Fig. 3D,E**). This gradient of change correlated well with distance from ablation site (rho=-0.7, *P*<2e-9). Recovery was limited in areas closest to the ablation (**Fig. 3D,E**).

### Outcome and Correlation to FA

Tremor significantly decreased and quality of life significantly increased following ablation (**Fig. 4A,B,C**). No correlation was found between FA values and tremor severity pre-ablation, but inverse correlations were found between the improvement in tremor and FA values in the thalamic motor area one day post-ablation in essential tremor (R=-0.62, *P*<0.003) and Parkinson’s disease (R=-0.52; *P*<0.03) patients. In essential tremor but not Parkinson’s disease, a similar inverse correlation was found between motor thalamus FA values pre-ablation and tremor improvement (**Fig. 4D**). To confirm that indeed the FA value pre-ablation can serve as a predictor for treatment outcome, an additional replication group of participants was recruited where only the first two time points were acquired. Test-retest demonstrated that both the discovery and replication groups independently show inverse correlation (R_discovery_=-0.51, *P*<0.02; R_replication_=-0.46, *P*<0.05). The inverse correlation remained when assessing tremor improvement at six months in the ET discovery group (n=20, R_discovery_=-0.49, *P*<0.04) but not the PD group (n=17, R=-0.035, *P*>0.1).

**Figure 4:**
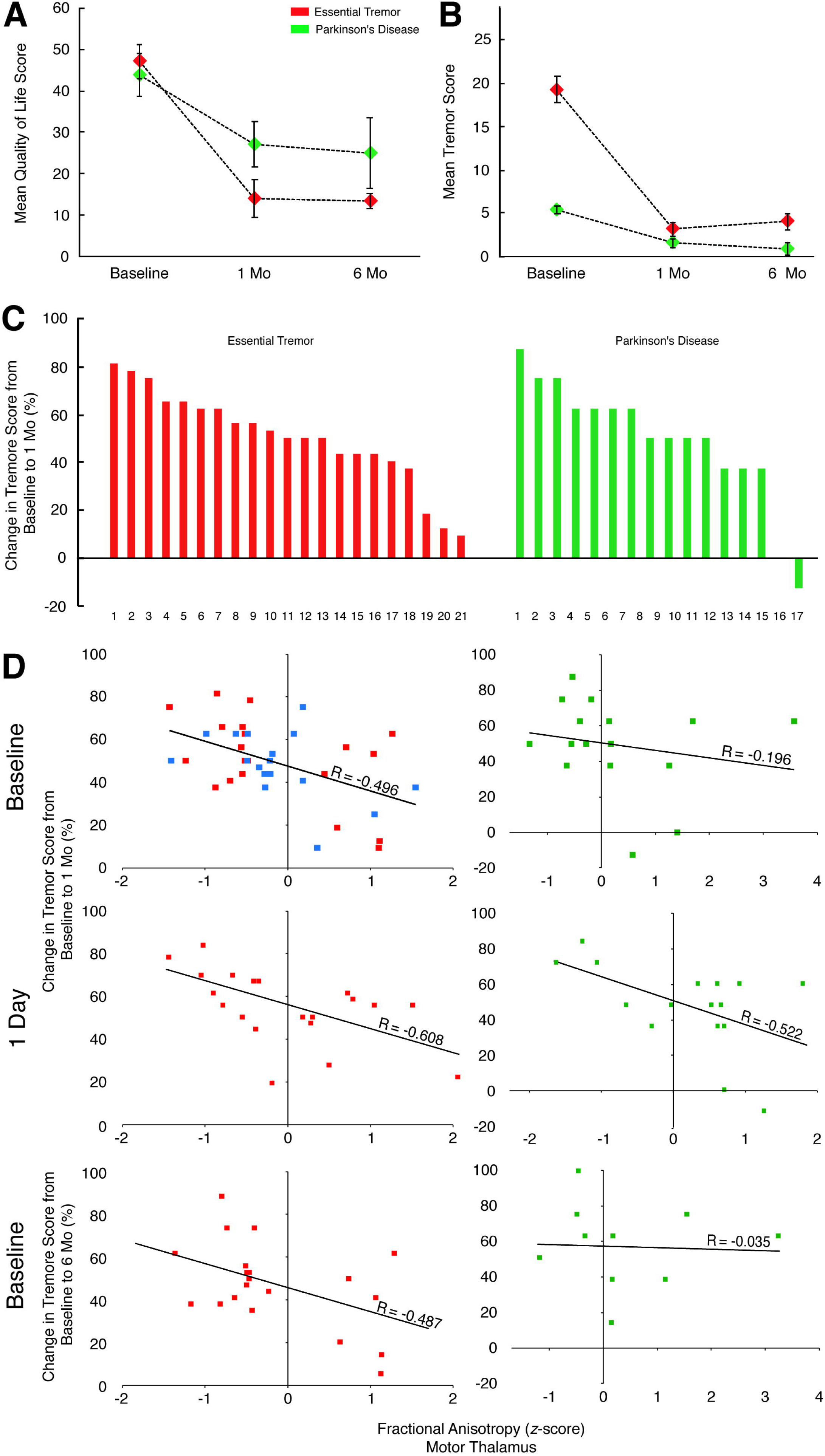
Tremor score evaluated before and after ablation reveals long-lasting improvement in both essential tremor (ET, *red*) and Parkinson’s disease (PD, *green*) study participants, and a correlation between measures of structural integrity before and after the ablation procedure and participant improvement. **(A)** Quality of life scores were calculated using the Quality of Life in Essential Tremor (QUEST) and Parkinson’s Disease Questionnaire (PDQ). (**B)** Tremor scores were evaluated using the Clinical Rating Scale for Tremor (CRST) and Unified Parkinson’s Disease Rating Scale (UPDRS). **(C)** Improvement of tremor one month post-ablation is shown for ET and PD study participants. **(D)** Correlation between tremor improvement and fractional anisotropy (FA) at the motor thalamus before treatment and one-day post-ablation demonstrate an inverse correlation between change in score one month after the ablation procedure and FA values for ET but not for PD. An additional group of seventeen ET study participants (*blue*) demonstrate the same effect of pre-treatment motor thalamus inverse correlation in a test-retest (see text for all details).

## Discussion

This study reports on white matter changes following unilateral focused ultrasound ablation to the motor thalamus. We observed a long-term focal FA decrease in the motor thalamus compared to a broader short-term decrease in areas along the DRTC tract. We examined the tracts between the ablated motor thalamus and different regions of interest in the motor system that are implicated in tremor generation. The tract to the red nucleus showed a consistent decline in tractography measures compared to other areas, which showed recovery to baseline values in the late follow-up scans. Examination of FA and MD changes throughout the DRTC tract indicated that local FA decreases co-occurring along with MD increases take place around the ablation site. Finally, for individuals with essential tremor but not Parkinson’s disease, we found an inverse correlation between the tremor score measured one month after the procedure and FA in the motor thalamus before ablation, implicating this region directly in the pathogenesis of these individuals. FA one day following ablation was correlated with tremor improvement in both patient groups.

### Diffusivity changes inside the Thalamus

In order to understand local thalamic changes, we assessed the lesion volume and extent of surrounding edema. FA values were evaluated in the ablated lesion, surrounding edema and different thalamic segments. As expected, volumes of the lesion and the edema increased immediately following ablation, with a gradual decrease in the late follow-up scans. The target of ablation within the motor thalamus, exhibited as a lesion, was the only region to show a permanent FA change. In contrast, FA in the edema area and in the sensory and motor portions of the thalamus recovered to baseline, while other parts of the thalamus were not affected by the ablation procedure, confirming the safety and accuracy of the procedure. Thus, the observed short-term changes in FA values may be partially or wholly attributed to the impact of edema on diffusivity.

Previous reports showed attenuation of contra-lateral tremor following stroke.^29–33^ The focal damage to the VIM induced by focused ultrasound ablation may share the same mechanism of tremor improvement occurring following stroke. A possible explanation is that the damage to VIM directly or indirectly (through feedback loops and white matter connections) affects a central oscillator located either within VIM itself or along its connections to the DRTC tract.

Short-term diffusivity changes were also seen inside the sensory thalamus, potentially explaining some of the adverse effects. Indeed, we found a correlation between the damage to the sensory thalamus reflected by an FA decrease and clinical side effects. A previous report showed the importance of the lesion location relative to white matter tracts crossing the thalamus.^34^ Our results replicate this prior observation and complement it by emphasizing that identification of sensory thalamus is beneficial in preventing side effects by refining MRgFUS treatment targeting.

### Ablation of the Motor Thalamus Results in Transient Changes in Most but Not All Fiber Pathways

A previous study,^17^ showed permanent FA decline in the tremor tract between the cerebellum and the cortex. Our study investigated the FA, MD, tract volume and number of tracts along the DRTC and known tremor areas. Contrary to this study, we found that FA and MD changes were restricted to the ablation site and its vicinity, and that all measures, including tract volume and number of tracts, returned to baseline between a few months to one-year post-ablation. This pattern of decline and recovery is consistent with the evolution of the edema at the ablation site and its surroundings. Thermal ablation induces edema that causes a decrease in FA, an increase in MD, and may affect tractography measures, similar to the effect of stroke.^34^

In contrast to other regions, the red nucleus demonstrated a consistent decline in tractography measures. The number of fibers and the volume of the white matter pathway between the red nucleus and the motor thalamus were not restored to baseline in the late follow-up scans suggesting that injury to this pathway may be chronic and related to tremor improvement. However, no correlation was found between diffusivity or tractography measures and tremor improvement. The red nucleus is comprised of magnocellular and parvocellular portions. Evolution from apes to humans and the transition from quadrupedalism to bipedalism caused a decrease in the size of the magnocellular portion and an increase in the size of the parvocellular portion. In primitive animals, the magnocellular portion affects motor function and locomotion using the rubro-spinal tracts. Hicks and Onodera^36^ suggested that pyramidal activity gradually replaced motor function of the magnocellular portion that remained in humans just as a backup for pyramidal motor activity. One possible mechanism is that in tremor disorders in which the motor control system is impaired, the function of the magnocellular portion and the rubro-spinal pathways are enhanced causing the tremor to emerge. A similar outcome has been demonstrated in the red nucleus following stroke.^37^ It is possible that damage to the connection between the thalamus and the red nucleus reduces this abnormal activity. The parvocellular portion affects tremor generation through the inferior olive and the Guillian-Mollaret triangle.^10^ Consequently, an alternative mechanism is that damage to the tract between the thalamus and the red nucleus changes the neural activity along this triangle, eventually affecting the dentate nucleus and its afferent fibers.

### Motor Thalamus Diffusivity Predicts Treatment Outcome in Essential Tremor but not in Parkinson’s Disease Patients

Our study tested the impact of focused ultrasound ablation in two distinct groups of tremor disorder patients: essential tremor and Parkinson’s disease. A previous study has already shown a difference in clinical improvement between these groups.^38^ We found no significant difference between the groups in terms of diffusion and tractography measures. Moreover, FA in the motor thalamus one day following ablation was inversely correlated with improvement in tremor score in both patient groups, suggesting that ablation to the motor thalamus impacts a tremor pathway common to both disorders. However, we identified that FA in the motor thalamus prior to the procedure is inversely correlated with an improvement in tremor score in essential tremor but not Parkinson’s disease patients, implicating the motor thalamus directly in the essential tremor pathogenesis.

The inverse correlation of pre-ablation FA and procedure outcome suggests that it may serve as a predictive measure of treatment success in essential tremor patients. FA represents normal fiber density,^39^ therefore, high FA in the motor thalamus may indicate normal white matter values having the ability to overcome the ablation by reconstructing the damaged white matter tracts. Alternatively, it is possible that intact white matter is less susceptible to be impacted directly by the ablation, therefore, residual pathological function remains following the treatment, while white matter impacted by the pathology in the motor thalamus is more vulnerable and subsequently damaged permanently by the ablation. Future studies with larger cohorts and in animal models of white matter pathology are needed to establish whether this imaging marker can be used for patient selection.

### Limitations

Experimental limitations include the dropout of patients who reached the late follow-up scans, reducing the signal-to-noise measurements relative to the early scans. In addition, we show temporal changes in tractography measures (number of tracts and tract volume) throughout the DRTC tract. These measures are calculated by a tractography algorithm that is known to have inherent limitations,^40^ thus may mis-estimate connectivity due to modeling errors. Further, the present study was limited in analyzing specific pathways. The cerebellar part of the cerebello-thalamo-cortical tract was not clearly demonstrated in the tractography analysis, possibly due to the limited number of directions acquired in the DTI and post-processing tractography algorithm limitations. Therefore, it was not possible to evaluate properly cerebellar changes in the white matter tracts.

## Conclusions

The clinical success of focused ultrasound ablation tremor therapy raises questions about the underlying mechanism in which thalamic nucleus lesioning reduces tremor. A deep understanding of white matter changes following focused ultrasound ablation could shed light on the mechanism of tremor and help tailor the procedure for the most appropriate patients. We found that DTI imaging provides evidence for long-term white matter damage in the ablation core, and in the tract connecting the thalamus and the red nucleus, suggesting it may play a role in tremor pathogenesis. It also shows a correlation between pre-ablation FA values in the motor thalamus and treatment success, thus implying its value as a possible biomarker for patient selection.

## Author Roles

1. Research project: A. Conception, B. Organization, C. Execution; 2. Statistical Analysis: A. Design, B. Execution, C. Review and Critique; 3. Manuscript Preparation: A. Writing of the first draft, B. Review and Critique

G.Z.: 1A-C, 2A-B, 3A

O.L.S: 1A-C, 2A-B, 3A

I.S: 1C, 3B

D.G: 3B

A.S: 1C, 3B

M.Z.: 3B

Y.A: 1A, 3B

A.E.: 1A-C, 3B

I.K: 1A-C, 2C, 3B

## Full financial disclosure for the previous 12 months

I.S. is a consultant for Actelion Pharmaceuticals Israel Ltd. and gave expert testimony for the Rehabilitation Department at the Israeli Ministry of Defense.

G.Z., O.L.S, D.G, A.S., M.Z., Y.A., A.E. and I.K. report no disclosures.

## Acknowledgements

This work was supported by the Israel Science Foundation 770/17 (I.K.), the Allen and Jewel Prince Center for Neurodegenerative Disorders of the Brain (I.K.) and the Adelis Foundation (I.K.).

The authors report no competing interests.

## Abbreviations

MRgFUS: magnetic resonance-guided focused ultrasound
VIM: ventral intermediate nucleus
DTI: diffusion tensor imaging
DRTC: dentato-rubro-thalamo-cortical
FA: fractional anisotropy
ICP: inferior cerebellar peduncle
MD: mean diffusivity
ADC: apparent diffusion coefficient
CRST: Clinical Rating Scale for Tremor
UPDRS: Unified Parkinson’s disease Rating Scale
TR: time repetition
TE: time echo
SPGR: spoiled gradient-recalled echo
FLAIR: fast fluid attenuated image recovery
MNI: Montreal Neurological Institute
FOD: fiber orientation distribution
ROIs: regions of interest
NOT: number of tracts
TV: tract volume

