## Supplementary Fig. 1 for "Structural integrity following focused ultrasound thalamotomy and its correlation with tremor relief"

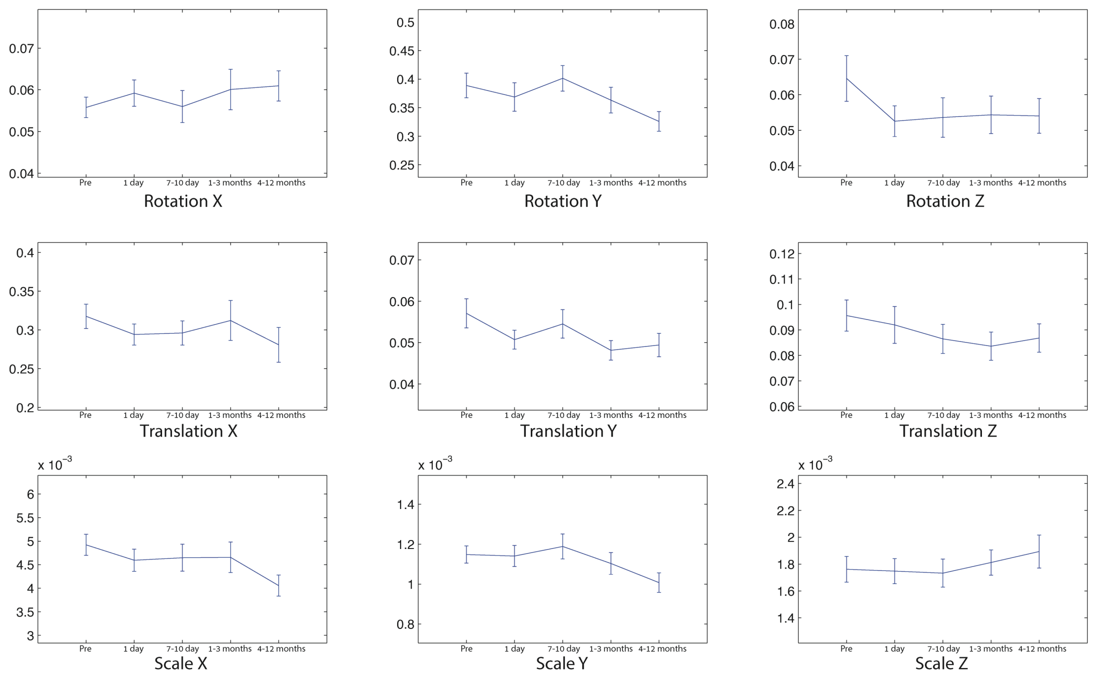


**Supplementary Figure 1:**

Motion values before correction for each time point (1, Baseline; 2, One Day; 3, 7–10 Days; 4, 1–3 Months; and 5, 6+ Months following ablation). Rotation (first row), translation (second row) and scaling (third row) are shown for x, y, and z axes (columns from left to right).
