## Supplementary Fig. 2 for "Structural integrity following focused ultrasound thalamotomy and its correlation with tremor relief"

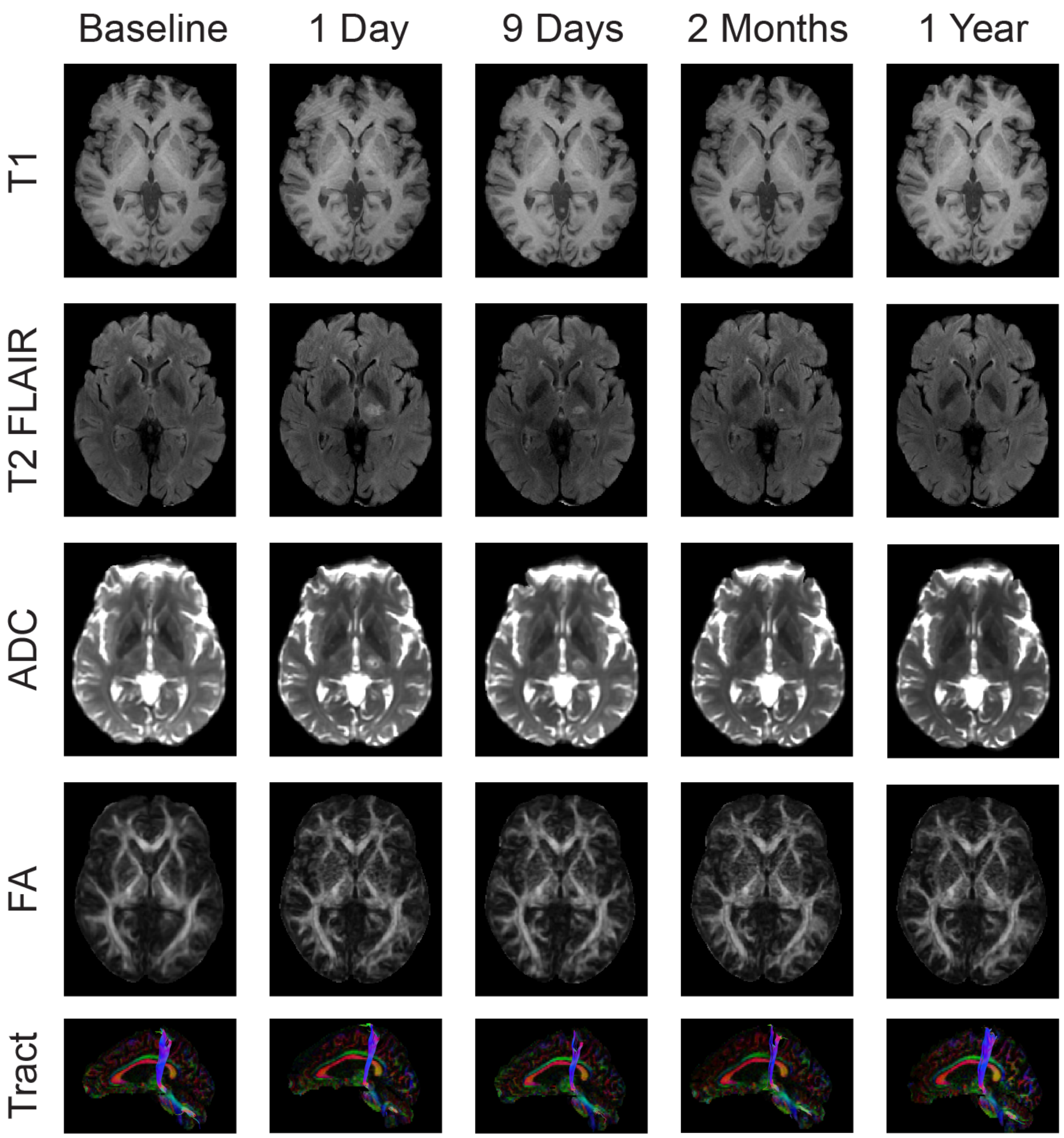


**Supplementary Figure 2:**

Temporal changes in the thalamic ablated lesion, surrounding edema and dentato-rubro-thalamo-cortical (DRTC) tract at all time points in one representative participant (65 years old, M, Parkinson's disease patient). T1, T2 fast fluid attenuated image recovery (FLAIR), apparent diffusion coefficient (ADC), fractional anisotropy (FA), and the calculated tract between the red nucleus and precentral gyrus. The lesion in the thalamus following the ablation procedure is expressed as an increase in ADC and a decrease in FA. Note that the intensity of T1 is decreased (first row). Signal changes in all MRI acquisition protocols are no longer visible in the last follow-up scan. Finally, the bottom row depicts the calculated tract between the red nucleus and precentral gyrus with similar recovery in the late follow-up scan.
