## Supplementary Table 1 for "Structural integrity following focused ultrasound thalamotomy and its correlation with tremor relief"

**Supplementary Table 1** Statistical differences (paired *t*-test) of volumes and FA at lesion and surrounding edema compared to pre-treatment (baseline) values

| | | 1 Day | 7–10 Days | 1–3 Mo | 6+ Mo |
|---|---|---|---|---|---|
| **Lesion Volume** | p-value | 4.93E-11 | 1.85E-07 | 5.94E-02 | 6.17E-03 |
| | df | 38 | 34 | 26 | 24 |
| | t-value | 7.50 | 4.46 | 1.36 | 1.31 |
| **Edema Volume** | p-value | 9.88E-10 | 6.98E-07 | 2.57E-02 | 1.72E-01 |
| | df | 37 | 33 | 25 | 23 |
| | t-value | 6.07 | 3.99 | 1.12 | -0.20 |
| **Lesion COG FA** | p-value | 1.69E-20 | 3.94E-16 | 1.89E-08 | 1.80E-06 |
| | df | 38 | 34 | 26 | 24 |
| | t-value | 18.00 | 14.17 | 7.68 | 5.98 |
| **Lesion entire extent FA** | p-value | 7.23E-22 | 1.83E-15 | 9.10E-09 | 1.24E-04 |
| | df | 38 | 34 | 26 | 24 |
| | t-value | 19.73 | 13.44 | 7.99 | 4.30 |
| **Edema COG FA** | p-value | 6.38E-10 | 7.25E-10 | 7.39E-05 | 7.63E-03 |
| | df | 37 | 33 | 25 | 23 |
| | t-value | 8.03 | 8.28 | 4.47 | 2.62 |
| **Edema entire extent FA** | p-value | 1.97E-19 | 3.00E-16 | 1.22E-07 | 3.96E-02 |
| | df | 37 | 33 | 25 | 23 |
| | t-value | 17.07 | 14.59 | 7.07 | 1.84 |

FA, fractional anisotropy; df, degrees of freedom; COG, center of gravity
