## Supplementary Table 2 for "Structural integrity following focused ultrasound thalamotomy and its correlation with tremor relief"

| **Supplementary Table 2** | Statistical differences (paired *t*-test) of FA in all thalamic regions compared to pre-treatment (baseline) values | | | | |
|---|---|---|---|---|---|
| | | **1 Day** | **7–10 Days** | **1–3 Mo** | **6+ Mo** |
| **Thalamus motor** | p-value | 1.69E-15 | 6.37E-13 | 1.11E-02 | 4.53E-02 |
| | df | 38 | 34 | 26 | 24 |
| | t-value | 12.49 | 10.73 | 2.39 | -1.73 |
| **Thalamus sensory** | p-value | 6.74E-13 | 1.71E-10 | 9.48E-03 | 3.87E-02 |
| | df | 38 | 34 | 26 | 24 |
| | t-value | 10.20 | 8.60 | 2.46 | -1.81 |
| **Thalamus frontal** | p-value | 4.65E-01 | 2.60E-01 | 4.68E-01 | 2.14E-03 |
| | df | 38 | 34 | 26 | 24 |
| | t-value | 0.09 | -0.64 | 0.08 | -3.09 |
| **Thalamus parietal** | p-value | 3.52E-01 | 3.99E-01 | 4.22E-01 | 6.25E-03 |
| | df | 38 | 34 | 26 | 24 |
| | t-value | 0.38 | -0.25 | -0.19 | -2.65 |
| **Thalamus occipital** | p-value | 4.40E-02 | 5.22E-02 | 4.40E-01 | 4.94E-02 |
| | df | 38 | 34 | 26 | 24 |
| | t-value | -1.73 | -1.64 | -0.15 | -1.68 |
| **Thalamus temporal** | p-value | 3.70E-02 | 5.14E-02 | 3.36E-01 | 2.71E-02 |
| | df | 38 | 34 | 26 | 24 |
| | t-value | -1.81 | -1.65 | 0.42 | -1.98 |

FA, fractional anisotropy

### Statistically significant following bonferroni correction
