## Supplementary Table 3 for "Structural integrity following focused ultrasound thalamotomy and its correlation with tremor relief"

Tractography values and statistical differences (non-parametric Wilcoxon signed-rank paired test) of NOT and TV at different ROIs compared to pre-treatment (baseline) values

| | | Baseline | 1 Day | 7–10 Days | 1–3 Mo | 6+ Mo |
|---|---|---|---|---|---|---|
| **SCP - NOT** | value | 378.84 | 255.24 | 320.00 | 381.31 | 342.43 |
| | p-value | | 0.035 | 0.203 | 0.218 | 0.798 |
| | z-value | | 2.105 | 1.273 | 1.232 | -0.256 |
| **SCP - TV** | value | 9831.13 | 7686.37 | 9721.83 | 9683.81 | 8558.42 |
| | p-value | | 0.008 | 0.163 | 0.118 | 0.716 |
| | z-value | | 2.633 | 1.393 | 1.562 | -0.363 |
| **CP - NOT** | value | 522.56 | 362.67 | 266.27 | 342.81 | 577.50 |
| | p-value | | 0.048 | 0.002 | 0.064 | 0.619 |
| | z-value | | 1.982 | 3.046 | 1.850 | -0.498 |
| **CP - TV** | value | 12181.11 | 9483.58 | 8407.88 | 10375.61 | 11946.86 |
| | p-value | | 0.100 | 0.015 | 0.149 | 0.904 |
| | z-value | | 1.647 | 2.421 | 1.441 | -0.121 |
| **RN - NOT** | value | 1663.31 | 1016.63 | 942.47 | 1180.79 | 1069.52 |
| | p-value | | 0.000 | 0.000 | 0.003 | 0.003 |
| | z-value | | 3.488 | 3.496 | 2.959 | 2.943 |
| **RN - TV** | value | 18961.76 | 14350.29 | 13588.57 | 14203.64 | 13600.46 |
| | p-value | | 0.001 | 0.041 | 0.003 | 0.130 |
| | z-value | | 3.183 | 2.043 | 2.959 | 1.514 |
| **PCG - NOT** | value | 387.85 | 240.38 | 278.18 | 376.79 | 348.37 |
| | p-value | | 0.039 | 0.025 | 0.881 | 0.526 |
| | z-value | | 2.065 | 2.234 | -0.149 | -0.635 |
| **PCG - TV** | value | 3431.67 | 2338.39 | 2148.66 | 3517.13 | 3019.82 |
| | p-value | | 0.008 | 0.000 | 0.970 | 0.520 |
| | z-value | | 2.670 | 3.652 | -0.037 | 0.643 |
| **PLIC PD - NOT** | value | 5903.06 | 4556.29 | 4355.93 | 6122.64 | 5863.50 |
| | p-value | | 0.004 | 0.001 | 0.534 | 0.966 |
| | z-value | | 2.911 | 3.237 | 0.622 | # |
| **PLIC PD - TV** | value | 45706.18 | 42145.88 | 38115.35 | 44856.69 | 43582.26 |
| | p-value | | 0.287 | 0.041 | 0.424 | 0.465 |
| | z-value | | 1.065 | 2.045 | 0.800 | # |
| **PLIC ET - NOT** | value | 6332.36 | 5238.55 | 4712.55 | 6385.22 | 6853.29 |
| | p-value | | 0.012 | 0.002 | 0.877 | 0.391 |
| | z-value | | 2.516 | 3.024 | 0.155 | # |
| **PLIC ET - TV** | value | 49132.48 | 44046.20 | 43432.21 | 52440.59 | 48001.17 |
| | p-value | | 0.168 | 0.296 | 0.756 | 1.000 |
| | z-value | | 1.380 | 1.045 | -0.310 | # |
| **SCR PD - NOT** | value | 3727.41 | 2538.18 | 2593.40 | 3972.55 | 3876.32 |
| | p-value | | 0.004 | 0.002 | 0.722 | 0.966 |
| | z-value | | 2.864 | 3.067 | 0.356 | # |
| **SCR PD - TV** | value | 30829.24 | 25812.82 | 24990.00 | 29539.46 | 29681.47 |
| | p-value | | 0.076 | 0.031 | 0.424 | 1.000 |
| | z-value | | 1.775 | 2.158 | 0.800 | # |
| **SCR ET - NOT** | value | 3446.36 | 2673.55 | 2612.85 | 3557.72 | 3695.71 |
| | p-value | | 0.00 | 0.02 | 0.84 | 0.30 |
| | z-value | | 3.003 | 2.427 | 0.207 | # |
| **SCR ET - TV** | value | 30296.02 | 26198.69 | 26828.87 | 33214.66 | 29198.36 |
| | p-value | | 0.020 | 0.079 | 0.717 | 0.952 |
| | z-value | | 2.321 | 1.755 | -0.362 | # |

### Number of subjects was insufficient for statistical testing
NOT, number of tracts; TV, tract volume; ROIs, regions of interest; SCP, superior corona radiate; CP, cerebral peduncle; RN, red nuclues; PCG, precentral gyrus; PLIC, posterior limb of internal capsule; SCR, superior corona radiata
